# A reusable model of pangenome selection informs optimal surveillance strategies over vaccine introductions

**DOI:** 10.64898/2025.12.18.695090

**Authors:** Leonie J. Lorenz, Joel Hellewell, Samuel T. Horsfield, Matthew J. Russell, Shrijana Shrestha, Andrew J. Pollard, Stephen D. Bentley, Stephanie W. Lo, Caroline Colijn, Nicholas J. Croucher, John A. Lees

## Abstract

**Background:** The human pathogen *Streptococcus pneumoniae* is a major cause of disease, including pneumonia and meningitis. The introduction of Pneumococcal Conjugate Vaccines (PCVs) initially reduced the burden of disease through a reduction of colonisation by vaccine-targeted serotypes. However, since PCVs only target a proportion of pneumococcal serotypes, they shift intraspecific competition, eventually allowing non-targeted types to ‘replace’ vaccine types. Understanding the host and pathogen factors causing replacement is important for future vaccine development. Mechanistic understanding of vaccine replacement dynamics is crucial for forecasting and optimisation of genomic surveillance strategies to evaluate realised vaccine effectiveness.

**Methods:** We developed a mathematical model of the genomic and demographic factors which explain vaccine replacement, used this model to replicate serotype-frequency changes, and investigated cost-effective genomic surveillance strategies. We extended a forward-time model based on the Wright-Fisher model, developing a user-friendly model framework that describes the post-vaccine dynamics of *S. pneumoniae* populations. Our model describes vaccine replacement as a function of vaccine impact, immigration of new strains, and negative frequency-dependent selection (NFDS) on the accessory genome content.

**Results:** We used our model to study vaccine replacement in newly sequenced genomic surveillance data from Nepal, and existing data from the US, and the UK, with distinct surveillance strategies. We showed that the model with NFDS better replicates replacement dynamics than a null model without NFDS, and that NFDS likely only acts on part of the *S. pneumoniae* accessory genome. We found consistent estimates for vaccination effectiveness across the different study locations and country-specific genes under NFDS, highlighting the importance of conducting genomic surveillance in each country of interest. By simulating data from the model, we showed that an optimal surveillance strategy prioritises per-sampling sample size over sampling frequency for small sampling budgets.

**Conclusions:** Our model can be used to predict vaccine replacement dynamics after PCV introduction, and can be easily reapplied to analyse new data from vaccine introductions or new regions. Our model is available in the R package STUBENTIGER (**Stu**dying **B**alancing **E**volution (**N**FDS) **T**o **I**nvestigate **Ge**nome **R**eplacement) on GitHub https://github.com/bacpop/Stubentiger.

## Background

*Streptococcus pneumoniae* is a human pathogen that causes pneumonia and meningitis, with the highest incidence of disease in children and elderly adults [1]. Pneumococcal Conjugate Vaccines (PCVs), first introduced in 2000 in the United States, successfully decreased the burden of disease and carriage of targeted serotypes [2, 3]. However, PCVs impact intraspecific competition, greatly reducing the fitness of the proportion of serotypes they affect and leading to the replacement of vaccine targeted serotypes by non-targeted serotypes [4, 5].

Traditionally, *S. pneumoniae* epidemiology has focused on serotypes alone, which are considered to be the main determinant of virulence [6] and are defined by the pneumococcal capsular polysaccharide (CPS). However, the variation of the *cps* locus only represents a small proportion of the genetic diversity. Whole genome sequencing has made it possible to investigate the full breadth of pneumococcal genomic diversity, including causes for invasiveness and antibiotic resistance, and allows the definition of closely-related lineages called global pneumococcal sequence clusters (GPSCs) [7]. While serotypes are defined only by the *cps* region, GPSCs are defined by the variation in the whole genome. Importantly, GPSCs and serotypes provide complementary information since many GPSCs contain multiple serotypes and *vice versa* [7]. Hence, serotype replacement can stem both from genetically similar serotypes from within the same GPSC or genetically different serotypes from other GPSCs [8].

The use of genomics has lead to major breakthroughs in microbial surveillance [9]. In the case of *S. pneumoniae*, our understanding of the pneumococcal population dynamics has been advanced by the Global Pneumococcal Sequencing (GPS) project. By September 2025, the GPS project had sequenced 25,997 pneumococcal genomes across 60 countries [10]. It is crucial to collect and leverage genomic surveillance data in order to understand the mechanisms of vaccine-type replacement, why certain serotypes are more successful in replacing vaccine-targeted serotypes, and ultimately, anticipate the future dynamics of *S. pneumoniae* populations. Additionally, the GPS project has informed optimisation of vaccine formulations and closed gaps in genomic surveillance data in countries with high burden of pneumococcal disease [11]. These countries tend to be low- and middle-income countries, where genomic surveillance faces stronger financial constraints [12], empirical use of antibiotics is higher and infrastructure of clinical microbiology labs is sparse, limiting the sample sizes and sampling frequency. Genomic surveillance therefore has the potential to guide the detection of underlying genetic mechanisms that drive ongoing replacement dynamics but requires careful planning in its rollout. A sampling strategy which was prospectively designed such that key questions about vaccine response could be answered would be a significant improvement over retrospectively attempting to answer these questions.

Negative frequency-dependent selection (NFDS) has been proposed as a major driver of vaccine replacement in *S. pneumoniae* [13, 14, 15]. NFDS is a type of selection that depends on the frequency of a given trait within a population: it is especially beneficial to have the trait when it is rare and less beneficial when the trait is common [16]. Examples of genes under NFDS in bacteria are antimicrobial resistance genes and bacteriocins.

Mathematical models using NFDS as an evolutionary process have been applied to describe vaccine replacement in *S. pneumoniae* populations [13, 14, 15]. These, and similar models, have helped to identify the strength of NFDS, vaccine impact, and immigration of new strains [17]. The same models can make predictions of future dynamics [18, 14]. However, we identified three key problems with previous modelling approaches. Firstly, previous models had been developed to work with specific datasets and were difficult to reuse by non-experts. This prevents public health bodies from reusing them in their own healthcare contexts, where the models could have the most impact. Secondly, former approaches had not compared the utility of genomic surveillance with serotyping or targeted sequencing data. It remained to be investigated whether the more resource-intensive collection of genomic surveillance data was necessary for effective post-vaccination surveillance or whether serotypes or sets of genes could be used as universal markers for NFDS, potentially allowing to transfer knowledge between countries and datasets. Thirdly, the impact of genomic surveillance sample size and frequency on a model’s ability to estimate epidemiological parameters of interest and predict future replacement dynamics remained unclear.

To solve the first problem of re-usability, we implemented a population dynamics model including NFDS, using the odin modelling framework [19]. By separating model code, genomic data input, and fitting procedures, our new model implementation is easy to adapt and re-use. We tested and compared using input of accessory gene data from state-of-the-art, reproducible bioinformatic tools, which we found superior to previous approaches. We applied our model to genomic surveillance datasets not only from the high-income countries US and UK, but also the lower-middle-income country Nepal. To solve the second problem of the utility of genomic surveillance data, we implemented different model versions that used serotyping or whole genome sequencing data. We built upon previous approaches of NFDS model comparison [13], which, combined with our likelihood-based approach, allowed us to perform rigorous model selection. Our likelihood-based approach also made it possible to test how many intermediate-frequency genes were under NFDS, which helped us to investigate the option of targeted sequencing approaches. To solve the third problem of sample size impact, we fitted the model to simulated data to investigate the effect of different surveillance strategies on parameter estimation and prediction, considering a scenario with limited genomic surveillance budget.

## Methods

Whole genome sequences from a carriage study in Kathmandu, Nepal, were obtained from the European Nucleotide Archive (ENA); the accession numbers can be found in Supplementary File 1. The isolates from Nepal were collected in 2009–2019 with 1881 carriage samples in total. Whole genome sequences and metadata for Massachusetts, US [20], and Southampton, UK [21, 22] were previously published. The sequences from the US were collected in Massachusetts in 2001, 2003, and 2007 with 616 sequences in total. The sequences from the UK were collected in Southampton in 2006–2012 with 672 sequences in total.

Where necessary, genomes were assembled using shovill 1.1.0 using default parameters [23]. Global pneumococcal sequence clusters (GPSCs) were computed using PopPUNK 2.6.0 [24] with the database for *S. pneumoniae* GPS v4 via poppunk assign, applying quality control (--run-qc), allowing maximally one distance of zero (--max-zero-dist 1), and merging a maximum of three clusters based on one single isolate (--max-merge 3).

We constructed a consistent pangenome that included all samples from our three datasets, which ensured that gene clusters were compatible across datasets. We first intended to compute separate pangenomes for each dataset and map the resulting gene clusters to each other using the easy-search function of MMseqs2 [25]. However, mapping two separate pangenomes with MMseqs2 gave a very different result than when computing a joint pangenome using pangenome tools, which we show an example of in Supplementary Figure S1. Hence, we decided to compute our consistent pangenome by giving the data from UK, US, and Nepal jointly to the pangenome tools in the following way: ggCaller 1.3.4 [26] was used for gene calling using the SPARC dataset for annotation [27, 28] and with the option --gene-finding-only. Bakta’s protein bulk annotation workflow (bakta proteins) [29] with default parameters was used for functional annotation and Panaroo v1.5.2 [30] was run with the options --refind-mode off --clean-mode moderate --family threshold 0.7 --min trailing support 2 --trailing recursive 0 --min edge support sv 2 --edge support threshold 0.0 --aligner none. Insertion sequence elements, which may generate paralogues within the dataset through inserting in multiple sites, were eliminated by removing genes with annotations containing any of ‘transposon’, ‘transposase’, ‘insertion sequence’, or ‘mobile’ (this removed approximately 2.8% of genes).

For the Massachusetts dataset, we compared the PopPunk output to sequence clusters created with BAPS [31] and clusters of orthologous genes generated with ggCaller and Panaroo to COGtriangles [32]. The comparison was achieved by fitting model versions on all four data processing combinations (BAPS + COGtriangles, BAPS + ggCaller/Panaroo, PopPUNK + COGtriangles, and PopPUNK + ggCaller/Panaroo) and comparing the resulting likelihoods.

We separated the pangenome matrix (output of Panaroo) for the three countries. Each of these pangenome matrices was then filtered for intermediate-frequency genes, i.e. we only kept gene entries that were present at 5–95 % frequency within each population. We made the assumption that the genetic variation (presence or absence of a gene) within the same GPSC is negligible. We therefore summarised the pangenome matrix, which states the presence/absence of genes for each isolate, for the GPSCs. For this, we took the median of all presences (i.e. if over 50 % of the isolates of a GPSC have a certain gene, the gene would be marked as present, otherwise as absent).

At the beginning of each simulation the GPSC-serotype combination counts are represented using a matrix, which we refer to as the start population. We generated the start population for the model by Poisson sampling from the GPSC-serotype counts from the first (UK and US) or first three (Nepal) time points. All GPSC-serotype combinations that do not appear in the first time point, but are present at later times, were initialised with 1.

In the model, GPSCs and serotypes immigrate into the population. The possible GPSC-serotype combinations are defined by the immigration matrix. We calculated the immigration matrix by assigning all GPSC-serotype combinations that appear at least once in the dataset equal probabilities of immigration (i.e. one over the number of GPSC-serotype combinations). The total population size of the model is constant and defined as 15, 000 isolates. The delta statistic, as introduced by Corander *et al.* (2017) [13], calculates the changes in gene frequencies over time. We calculated it in the same way as described in their paper, by calculating the changes in gene frequencies across loci *l* of the first time point (UK and US) or first three (Nepal) *e*, which are assumed to be at equilibrium, to the gene frequencies of the last time point *f* . Genes that change the least in frequency have a low delta statistic value, while genes that change more have a higher value. For gene *l* the delta statistic is defined as 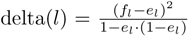. For convenience, we then ranked the delta statistic values for all genes. We calculated the ranked statistic independently for the US, UK, and Nepal.

Other inputs to the model include a vector that defines which serotypes are affected by the vaccine and when the vaccine rollout started, both of which can be typically found in the literature. The model also needs the information of how many generations of *S. pneumoniae* to simulate between each data sampling time point. We generally assumed that there was one *S. pneumoniae* generation per month, meaning that there would be 12 generations per year. This should be given in the form of *dt* = 1*/*generations between sampling time points, e.g. *dt* = 1*/*12.

Our population dynamics model was based on the model from Corander *et al.* (2017) [13], which is an extension of the Wright-Fisher model to include immigration, and selection due to vaccination and NFDS. While similar, our model is formulated as a compartmental model rather than an individual-based model. We therefore assume that no novel genotypes can arise during simulation, as this would lead to a variable-dimensional state space, which makes inference numerically challenging.

All model versions of our population dynamics model expect the following data-informed inputs described in the section above: the GPSC-summarised pangenome matrix, start population, immigration matrix, delta ranking, vaccine vector, vaccine rollout time point, and *dt*.

In the following, we will describe the *partial NFDS, strain-serotype* model, which has four parameters that can be fitted to data and considers some but not all genes to be under NFDS (*partial NFDS* ). Other model versions used for comparison are described below. The *strain-serotype* model represents the population at time *t* as the matrix *P^t^*, with rows representing distinct strains (GPSCs) *i* = 1*, . .., n* and columns representing distinct serotypes *j* = 1*, .. ., l*. Generations of *P^t^* are non-overlapping, and the subsequent generation *P^t^*^+1^ consists of the offspring population *B^t^* and immigrating strains *M^t^* calculated from the current generation 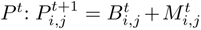

The immigrating strains *M^t^* are binomially distributed with success probabilities defined by the immigration matrix *m_i,j_*, which represents all observed strain-serotype pairs in the data, weighted with equal probabilities, and number of experiments defined by the immigration number *m̂ ^t^*. *m̂ ^t^* is binomially distributed with number of experiments determined by the carrying capacity *K* and probability of success determined by the immigration parameter *m*, which is fitted to data. The carrying capacity *K* is the constant population size of the model population and can be chosen by the user.

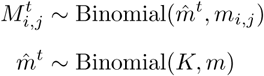

The offspring population *B^t^* depends on the carrying capacity *K*, the non-immigrating force 1 − *m*, and the offspring matrix *p_i,j_* of the population *P^t^*. The offspring matrix *p_i,j_* represents the probabilities of transmission to the next generation, depending on the NFDS term 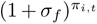 , the number of individuals in each compartment in the parent generation 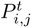, and the vaccination effect 1 − [*v*_time_(*t*) *· v*_type_(*j*) *· v*].

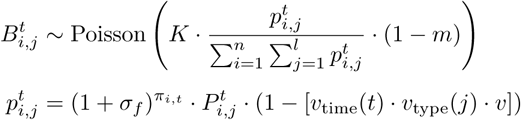

The NFDS term 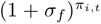 consists of the parameter for the strength of NFDS *σ_f_* and the deviation from equilibrium gene frequencies for strain *i*, *π_i,t_*. *π_i,t_* is calculated by subtracting the current gene frequencies *f* from the equilibrium gene frequencies *e*, for all genes *G_i_* that are found in strain *i* and that are in the set of genes under NFDS *S*. *S* is determined by ordering intermediate-frequency genes according to the delta statistic described above and making a cut-off at the proportion defined by parameter *prop_f_* .

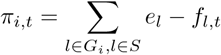

The vaccination effect 1 − [*v*_time_(*t*) *· v*_type_(*j*) *· v*] consists of function *v*_time_(*t*), which returns 1 if the model time is at a post-vaccine time point; the function *v*_type_(*j*), which returns 1 if serotype *j* is a vaccine type; and the vaccination effectiveness parameter *v*, which can be fitted to data.

In summary, the parameters that can be fitted to data are the vaccination effectiveness parameter *v*, the NFDS strength *σ_f_* , the proportion of genes under NFDS *prop_f_* , and the immigration rate *m*. The *partial NFDS, strain-serotype* model is implemented in our R package STUBENTIGER (**Stu**dying **B**alancing **E**volution (**N**FDS) **T**o **I**nvestigate **Ge**nome **R**eplacement), which includes functions for simulating and fitting models, as well as an example dataset. It can be installed from GitHub https://github.com/bacpop/Stubentiger.

Other model versions are specific variations of the *partial NFDS, strain-serotype* model described above.

- In the *no-NFDS, strain-serotype* model, *σ_f_*is fixed to 0 and *prop_f_* is fixed to 1, as described in Corander *et al.* [13]. This is a null model that has no notion of NFDS.
- In the *general-NFDS, strain-serotype* model, *prop_f_* is fixed to 1, hence all intermediate-frequency genes are under NFDS, as described in Corander *et al.* [13].
- In the *varying-NFDS, strain-serotype* model, which introduces two gene compartments; one under strong NFDS and one under weak NFDS, the strength of which is denoted by the additional parameter *σ_w_*, as described in Corander *et al.* [13].
- The *strain* model only contains frequencies of GPSCs and has no concept of serotypes. In this model, a GPSC is affected by a vaccine if the majority of serotypes within a GPSC are vaccine types.
- The *serotype* model only contains frequencies of serotypes and has no concept of GPSCs. For the data-informed inputs, genetic information is summarised per serotype and the likelihood compares the model outputs to serotpe counts instead of GPSC counts.

Data analysis and plotting was done using R 4.4.1 and RStudio 2024.09.0+375. Our population dynamics model was implemented using the R packages odin.dust (version 0.3.13), odin (version 1.5.11), dust (version 0.15.3), and mcstate (version 0.9.22) [19]. The code can be found on GitHub https://github.com/bacpop/NFDS_Model.

The models were fitted using Markov Chain Monte Carlo (MCMC) using the R package mcstate [19]. For MCMC, we defined a multinomial likelihood. We compared our model to GPSC counts for all sampling time points. Our multinomial likelihood takes these counts and compares them to the model values by summing over all serotypes of a GPSC.

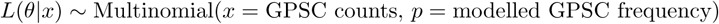

Because *σ_f_* and *m* tend to be close to zero, we fitted the logarithmic value of these parameters, which makes it easier for MCMC to sample. We defined uniform priors on [0, 1] (which for the two transformed parameters *σ_f_* and *m* resulted in 1*/x* priors after transformation).

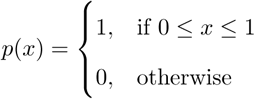

We ran four chains of the MCMC in deterministic mode with a deterministic particle filter (which takes the mean of the probability distributions in the model), first for 2000 steps and then for an additional 20, 000 steps. For Nepal, fitting took longer and we therefore ran the chains for 1000 steps and then 10, 000 steps instead. We applied the Chi-Squared distribution function (pchisq) in R to execute likelihood-ratio tests for model comparisons, applying the Bonferroni correction to compensate for multiple testing. We also calculated the Bayesian Information criterion (BIC) to compare model likelihoods.

To compare the GPSC- and serotype-frequency changes of our model and the data, we assumed that the data was generated through multinomial draws from the underlying population, to allow us to represent uncertainty of the data due to sample size. We then computed 95 % confidence intervals by calculating the 0.025 and 0.975 quantiles.

For analysing whether the same genes were found to be under NFDS in different datasets, it was crucial to have the consistent pangenome we described above. However, while the gene clusters were consistent, there were differences in gene frequencies between the countries. To be able to compare the sets of genes under NFDS, we only included genes that were at intermediate frequency in all three countries.

As an alternative approach to the delta statistic described above, we employed a genetic algorithm to find genes under NFDS. We used the genetic algorithm implemented in the R Package GA [33]. The genetic algorithm started with a variety of vectors representing all intermediate-frequency genes, which could be affected by NFDS (one) or not (zero) (’the population’). The algorithm then tried to find the best combination of ones and zeros by changing individual values (’mutation’), combining different zero-one vectors (’recombination’), and choosing the best candidates according to the likelihood described above (’selection’). The selection was informed by feeding the chosen NFDS vector into a version of the *partial NFDS, strain-serotype* model that takes an NFDS vector as input and returns the likelihood. The parameters of the model (vaccination effectiveness, NFDS strength, proportion of genes under NFDS, and immigration rate) were fixed.

To test whether the overlap between genes under NFDS of different countries or methods was larger than to be expected at random, we calculated the expected overlap between two (or multiple) gene sets.

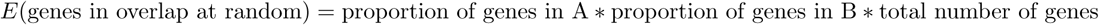

To generate data for the simulation study of genomic surveillance strategies, we simulated the *partial NFDS, strain-serotype* model 20 years forward in deterministic mode, using data inputs and parameter estimates from Massachusetts, US. We then sub-sampled this synthetic dataset to 0.1%, 0.25%, 0.5%, 1%, 2.5%, 5%, 10%, 25% and for annual, biennial, triennial, and quadrennial sampling. We then fitted the model as described above and with three independent replicates. After fitting the model, we sampled parameter estimates from the posterior distribution, simulated the model forward with these parameters and computed a cumulative mean squared error (MSE) as compared to the original simulated data across all 20 simulation years.

## Results

Our model describes the population dynamics of *S. pneumoniae* as a function of vaccination, negative frequency-dependent selection (NFDS), and immigration of global pneumococcal sequence clusters (GPSCs) and serotypes. Model outputs are serotype/GPSC density for each modelled time point. Our model is designed to take genomic surveillance carriage data, spanning at least one vaccine introduction, as input, to which the model can be fitted to infer parameters (Figure 1A). An overview of the data, input, model structure, and output is shown in Figure 1.

**Figure 1:**
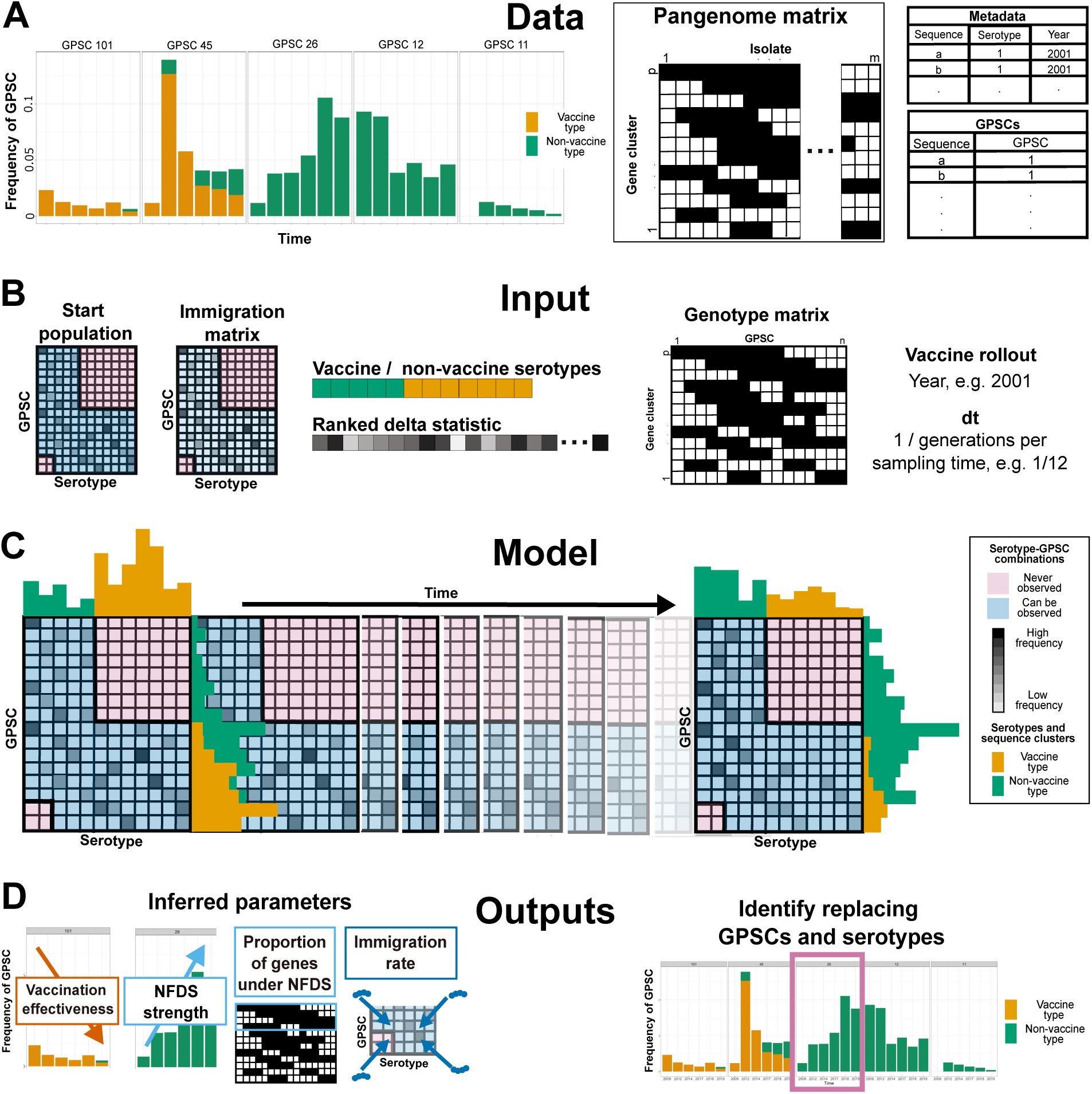
*S. pneumoniae* surveillance data, input, model structure, and model outputs. A) The Data panel shows illustrative frequencies of *S. pneumoniae* GPSCs, a pangenome matrix, a metadata and GPSC data table. B) The Input panel shows the data-informed inputs for the model: the start population, the immigration matrix, the vaccine-type vector, the ranked delta statistic, the genotype matrix, the year of the vaccine rollout, and model generations per sampling time. C) The Model panel shows the two-dimensional Wright-Fisher model structure, representing GPSCs and serotypes. Marginal frequencies represent overall serotype and GPSC frequencies, respectively, and can be compared across time points. D) The Outputs panel illustrates the four parameters estimated by the *partial NFDS, strain-serotype* model and that replacing GPSCs or serotypes can be identified using the model.

The population dynamics model is a compartmental Wright-Fisher model with selection. The compartmental structure of the model is two-dimensional, one axis represents GPSCs and the other axis represents serotypes (Figure 1C). The code outputs the modelled GPSC-serotype frequencies for any time step, as well as the GPSC or serotype frequencies separately (Figure 1D). In our model the number of compartments is fixed, which makes it less flexible than an equivalent individual-based model but faster to simulate from and fit.

The model is implemented in the domain-specific language odin [19]. Odin has a very similar syntax to R and can be used in RStudio, making it more easily extensible than bespoke compiled code. Furthermore, model training and inference is fast because model code is transpiled to C++, leveraging its runtime advantages. In odin, the model code is separate from the model simulation and fitting, which makes it easy to share models and run them separately from these components. Odin also allows the user to switch between stochastic and deterministic model simulation, which means users can switch between more realistic simulations and faster, deterministic ones.

We designed our model to be used with standard genomic surveillance carriage data of *S. pneumoniae*, which is collected by taking nasal swabs from healthy children across a number of years, followed by culture and whole genome sequencing. To observe dynamics away from endemic equilibrium, samples should ideally be taken before and after vaccine introduction. It is necessary that the data includes the time of sampling. The following data processing steps need to be taken before the data can be given to the model:

1. Sequences should be grouped into GPSCs using PopPUNK [24], which represent genetically similar groups and allow cross-dataset comparisons because GPSCs are consistent.
2. The serotype of each sequence must be determined, which can be done in a laboratory or using bioinformatic tools, like SeroBA(v2.0) [34]. Both determining GPSCs and serotypes can be easily done using the GPS Pipeline [35].
3. The accessory genome must be determined. A good option is ggCaller [26] for calling genes and Panaroo
4. [30] for building a pangenome graph. To input the pangenome matrix into the model, gene presence/absence must be summarised per GPSC since we assumed that within-GPSC variation is negligible. ggCaller and Panaroo are reliable for identifying orthologous genes, which is crucial for correctly estimating multilocus NFDS. We showed that the model fits using ggCaller and GPSCs (from PopPUNK) are equivalent or better than fits based on former state-of-the-art COGtriangles [32] and BAPS [31], which are no longer maintained or need more manual curation, in Supplementary Table S1 and Figure S2.

The data then needs to be converted to reflect the input format for the model. This is illustrated in Figure 1B and described in the Methods section.

The model can be fitted to surveillance data with Markov Chain Monte Carlo (MCMC) using mcstate [19]. MCMC estimates a posterior distribution for each parameter value, giving estimates and uncertainty, and allowing for incorporation of priors. Model comparison can be achieved using the model likelihood.

The outputs of our model can help to understand the ongoing vaccine replacement dynamics. They reflect the population dynamics over time and can help to identify replacing GPSCs and serotypes. Similar models have been used to explore the impact of different vaccine designs [18]. The model outputs also include inferred parameter estimates: first, the vaccination effectiveness *v*, which consists of vaccine effectiveness and the coverage of the vaccine rollout, and describes how much the fitness of vaccine types is reduced by the vaccine; second, the NFDS strength *σ_f_* , which measures the impact of NFDS on the genes which are affected by NFDS, and which confers a frequency-dependent fitness effect on carriers of these genes; third, the proportion of genes under NFDS *prop_f_* , which determines how many of the intermediate-frequency genes are affected by NFDS; and fourth, the immigration parameter *m*, which describes how many members of an external population enter the internal population per generation. These four parameters also exist in the NFDS model by Corander *et al.* (2017) but their best-fitting model includes an additional parameter, the “weaker NFDS strength” [13]. This parameter did not improve the fit of our population dynamics model, which we show below. Because the population dynamics model considers NFDS to have an effect on some but not necessarily all intermediate-frequency genes, we will refer to this model as the *partial NFDS, strain-serotype* model.

We studied vaccine replacement dynamics in genomic surveillance data from the US, the UK, and Nepal. These three datasets represent different surveillance strategies and introduction schedules of different PCV formulations. The dataset from the US covers the introduction of the first vaccine PCV7 (affecting serotypes 4, 6B, 9V, 14, 18C, 19F, and 23F), and the sampling was done with increasing sample sizes and consistently every three years (Figure 2A, right). The dataset from the UK covers the introduction of both PCV7 and PCV13 (affecting serotypes 1, 3, 5, 6A, 7F, and 19A in addition to those in PCV7), and sampling was done consistently every year (Figure 2A, middle). The dataset from Nepal covers the introduction of PCV10 (affecting serotypes 1, 5, and 7F in addition to those in PCV7, with cross immunity for 6A [36]), and sampling was done less consistently, with varying sample size and sampling frequency but an overall larger sample size and for a longer time (Figure 2A, left).

**Figure 2:**
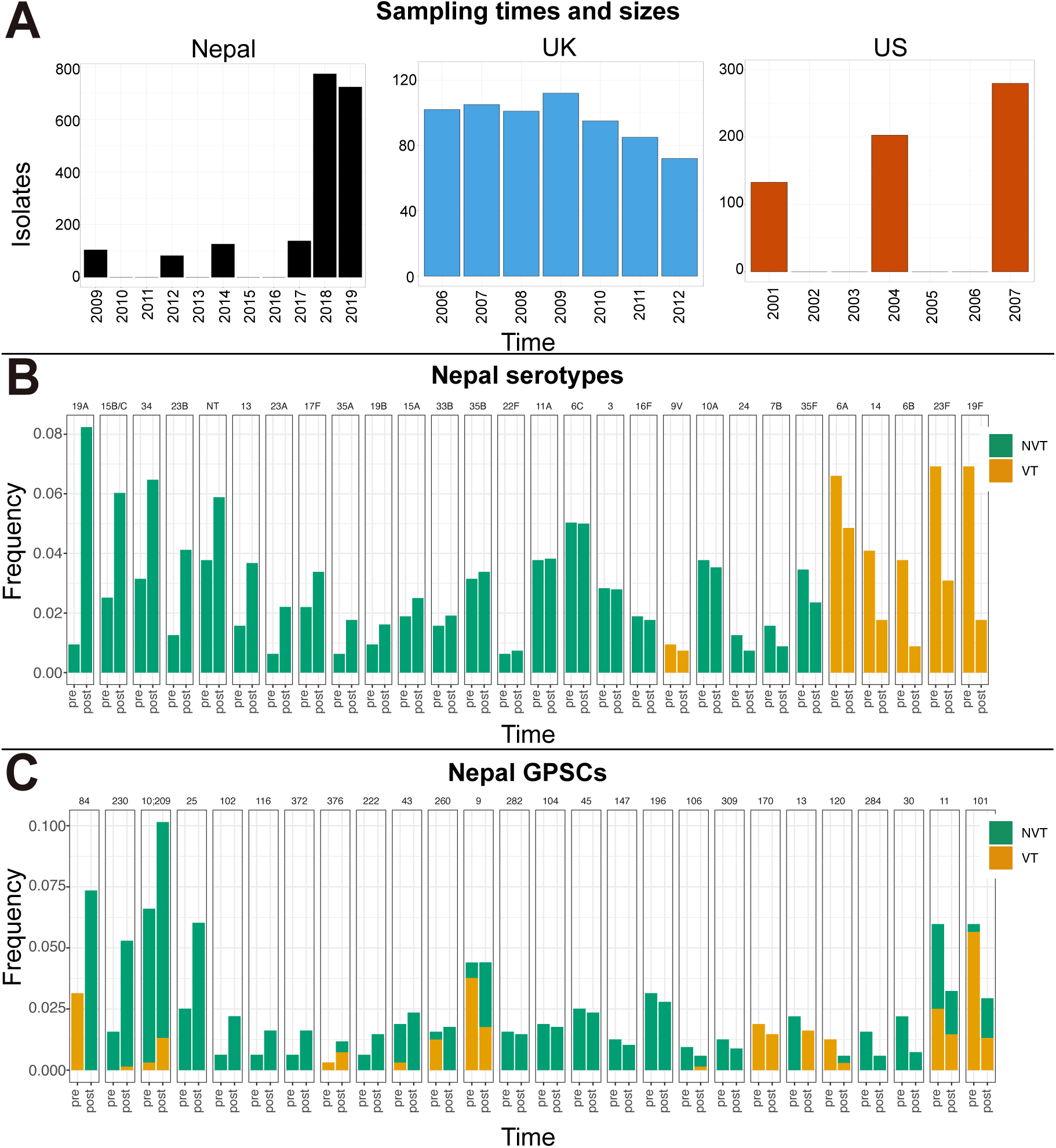
Sample sizes for Nepal, US and UK, and pre- and post-vaccine serotype and GPSC frequencies for Nepal. A) Sample sizes and sampling years for Nepal, UK, and US. B) Serotype and C) GPSC frequencies for pre-vaccination time points (mean of 2009, 2012, and 2014) and last post-vaccination time point (2019), sorted from largest positive change to largest negative change. Non-vaccine types (NVTs) shown in green and vaccine types (VTs) shown in orange.

Figure 2B and 2C show Nepal’s pre- and post-vaccine frequencies of serotypes and GPSCs, respectively, ordered by frequency changes. Panel B shows that vaccine types reduce in frequency, while non-vaccine types respond variably: some increase in frequency, some are stable. Panel C shows that the GPSC frequencies tend to be more stable. GPSCs containing vaccine types tend to reduce in frequency but can be partially rescued by non-vaccine types in the same GPSC.

We fitted the *partial NFDS, strain-serotype* model to each of these genomic surveillance datasets separately. Figure 3A to F compare the changes in serotype frequencies with 95 % confidence intervals from the data and the *partial NFDS, strain-serotype* model. The plots are split into non-vaccine types (NVTs), A, C, E, and vaccine types (VTs), B, D, F, for Nepal, US, and UK, respectively. When data confidence intervals overlap zero, frequency changes are not significant. Overlapping confidence intervals between model and data can be interpreted as agreement of model and data. All model confidence intervals for all countries overlap with the confidence intervals of the data. The model correctly describes whether the mean frequency of a serotype has increased or decreased for almost all serotypes. Interestingly, for Nepal, serotype 6*A*, which is affected by cross-immunity [36], but is not included in PCV10, shows a weaker response to the vaccine in the data. Our model does not capture this effect as we included 6*A* as a standard VT. Similar model fits to US and UK datasets have been previously reported in Corander *et al.* (2017) [13], which we used to validate our model formulation and fits. A comparison of model and data GPSC-frequency changes is shown in Supplementary Figure S3.

**Figure 3:**
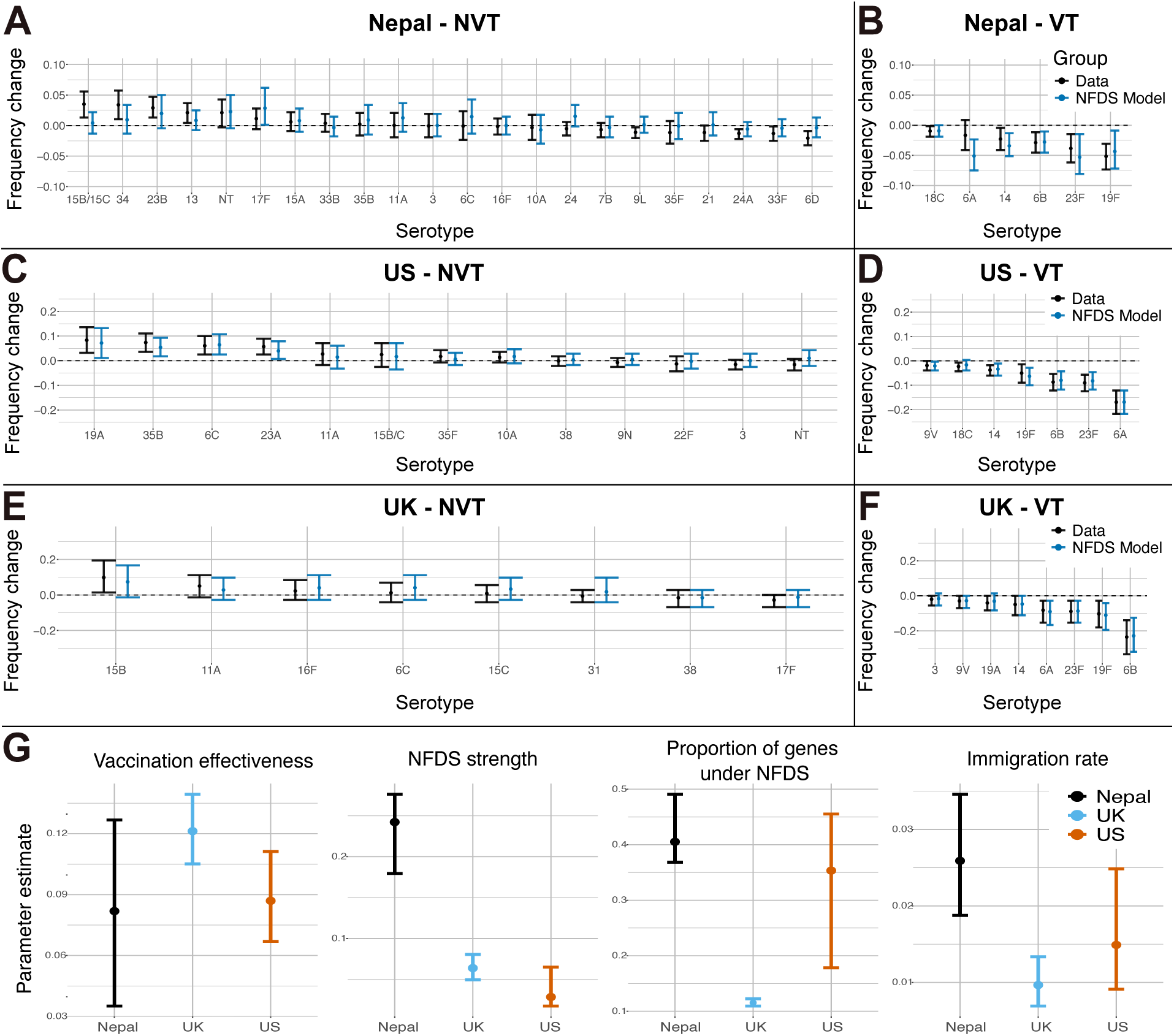
Comparison of *partial NFDS, strain-serotype* model to data and parameter estimate comparison across countries. A) - F) Comparison of serotype frequency changes in the data and forward simulation of *partial NFDS, strain-serotype* model with mean and 95% confidence intervals. Only serotypes with a frequency of larger than one percent in the pre-vaccine sample are shown. A) Nepal non-vaccine types (NVTs), B) Nepal vaccine types (VTs), C) US NVTs, D) US VTs, E) UK NVTs, F) UK VTs. G) Parameter estimates for vaccination effectiveness, NFDS strength, proportion of NFDS, and immigation rate with mean and 95% credible intervals for Nepal, UK, and US.

Figure 3G compares the mean parameter estimates and 95 % credible intervals of the *partial NFDS, strain-serotype* model across the three datasets from Nepal, US, and UK. The vaccination effectiveness is consistent across all locations and the mean estimate ranges between 0.082 and 0.121. The immigration rate is relatively consistent as well, with its means ranging between 0.0097 and 0.0259. The proportion of genes under NFDS is similar in the US and Nepal but lower in the UK. The NFDS strength is similar in the US and the UK but appears to be stronger in Nepal. This is potentially be driven by the within-GPSC replacement in Nepal, which was shown in Figure 2C.

Whether all intermediate-frequency genes are affected by NFDS has both ecological and epidemiological implications. It is ecologically relevant because intermediate-frequency genes are expected to sweep or go extinct when there is only drift and directional selection [37]. Investigating whether NFDS alone can explain stable pangenomes is therefore informative. Concerning epidemiology, NFDS might help define gene markers if it only affects a subset of all intermediate-frequency genes, facilitating to monitor GPSC frequencies or to determine gene function and design of future vaccines.

To determine whether NFDS affected all intermediate-frequency genes, we implemented three additional models. The first model was *no-NFDS, strain-serotype* model, which is a null model without the notion of NFDS. It only contains the two parameters of vaccination effectiveness *v* and immigration rate *m*. The second model was the *general-NFDS, strain-serotype* model, which has one additional parameter compared to the *no-NFDS, strain- serotype* model, the NFDS strength *σ_f_* , which affects all intermediate-frequency genes. Unlike the *partial NFDS, strain-serotype* model, the *general-NFDS, strain-serotype* model does not contain a parameter for the proportion of genes under NFDS; all intermediate-frequency genes are assumed to be affected equally by NFDS. The third model, the *varying-NFDS, strain-serotype* model, has an additional parameter compared to the *partial NFDS, strain-serotype* model, the weaker NFDS strength *σ_w_*, which affects all intermediate-frequency genes that are not affected by the other NFDS parameter, as determined by the proportion of genes under NFDS parameter. These three models had previously been implemented by Corander *et al.* (2017) and the authors found the *varying-NFDS, strain-serotype* model to produce the best fits [13].

We fitted all four models to the three datasets and compared the quality of the fits. Both likelihood-ratio tests and Bayesian Information Criterion (BIC) allow a statistical comparison of our models’ likelihoods while considering the number of free parameters (degrees of freedom). The likelihood-ratio test is significant when additional parameters are significantly improving the likelihood and the BIC is lowest for the model version that best explains the data. The *partial NFDS, strain-serotype* model was significantly better than the *no-NFDS, strain-serotype* model and the *general-NFDS, strain-serotype* model, and the *varying-NFDS, strain-serotype* model was not significantly better than the *partial NFDS, strain-serotype* model. The BICs and the significance of the likelihood-ratio test can be found in Table 1.

**Table 1:**
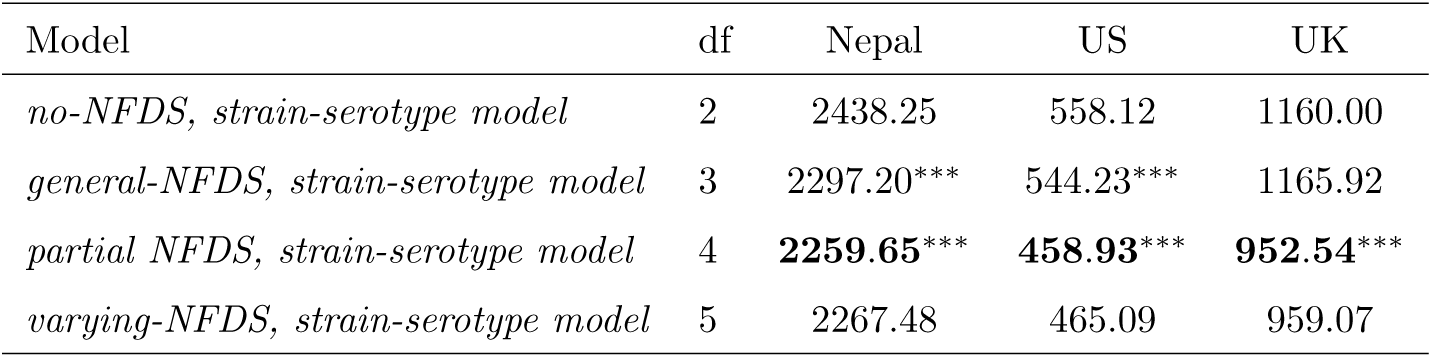
BIC and likelihood-ratio test on fits of different model versions. Model versions with degrees of freedom (df) are shown with Bayesian Information Criterion (BIC) and whether they are significantly better (*<* 0.0001, ***) than the simpler model versions (with less dfs) according to likelihood-ratio test. The best model version for each country is shown in bold.

| Model | df | Nepal | US | UK |
| --- | --- | --- | --- | --- |
| *no-NFDS, strain-serotype model* | 2 | 2438.25 | 558.12 | 1160.00 |
| *general-NFDS, strain-serotype model* | 3 | 2297.20*** | 544.23*** | 1165.92 |
| *partial NFDS, strain-serotype model* | 4 | **2259.65\*\*\*** | **458.93\*\*\*** | **952.54\*\*\*** |
| *varying-NFDS, strain-serotype model* | 5 | 2267.48 | 465.09 | 959.07 |

The result that the *partial NFDS, strain-serotype* model was consistently the best model version demonstrates two things: First, NFDS significantly increases the quality of the fit, suggesting that NFDS plays a role in vaccine replacement dynamics. Second, not all the intermediate-frequency genes identified by Panaroo show signs of being affected by NFDS. While NFDS is thought to strongly contribute to the stability of the accessory genome, it might not be the only factor maintaining the accessory genome. Factors like balanced rates of gene gain and loss or the occupation of different ecological niches might contribute to a stable accessory genome as well [37].

Whole genome surveillance is still costly and not available for many past vaccine introductions, which poses the question of whether other data types from surveillance, such as serotyping or targeted sequencing approaches, carry enough information to successfully deploy our population dynamics model.

We first investigated a scenario where only serotyping data was collected but some information about the genetic composition was available (e.g. from other countries or previous studies). To emulate this scenario, we implemented a model version that only contained the frequencies of serotypes (the *serotype model* ), summarised the pangenome matrix by serotypes and, in the likelihood, compared the model outputs to serotype counts from the data. The *serotype model* does not fit well to the data, as shown in Supplementary Figure S4. Hence, serotype information alone is not sufficient for deploying our population dynamics model.

We then intended to define a set of genes under NFDS that was consistent across different *S. pneumoniae* populations. Such a gene set could aid vaccine replacement studies because future models would not rely on computing the delta statistics for identifying genes under NFDS, which uses pre- and post-vaccine gene frequencies. This would increase the model’s ability to make predictions. A possible implication would also be to use targeted sequencing as a surveillance strategy, by focusing sequencing efforts primarily on NFDS genes and thereby reducing costs. Potentially, such a gene set could also inform future vaccine development because vaccine replacement dynamics could be understood before the introduction of a new vaccine and proteins that are affected by NFDS could possibly be targeted by serotype-protein vaccines.

We applied a genetic algorithm to identify such a consistent NFDS gene set. Genetic algorithms are optimisers that are inspired by natural selection. They are typically applied to a ‘population’ of vectors, in which they generate diversity through changing values (‘mutation’) and combining vectors (‘recombination’) and then keep the best vectors according to some fitness function (‘selection’). This process is repeated multiple times (‘generations’). When applying the genetic algorithm, we used the same likelihood as in the other (delta statistic-based) *strain-serotype* models and hence were able to compare these likelihoods directly. Overall, the genetic algorithm performed variably compared to the the delta statistic-based approach. While the genetic algorithm resulted in a better likelihood than the *no-NFDS, strain-serotype* model and *general NFDS, strain-serotype* model but not than the *partial NFDS, strain-serotype* and *varying NFDS, strain-serotype* model for the US, it resulted in worse likelihoods than any of the delta statistic-based models for the UK, and did better than all of the delta statistic-based models for Nepal, see Figure 4A.

**Figure 4:**
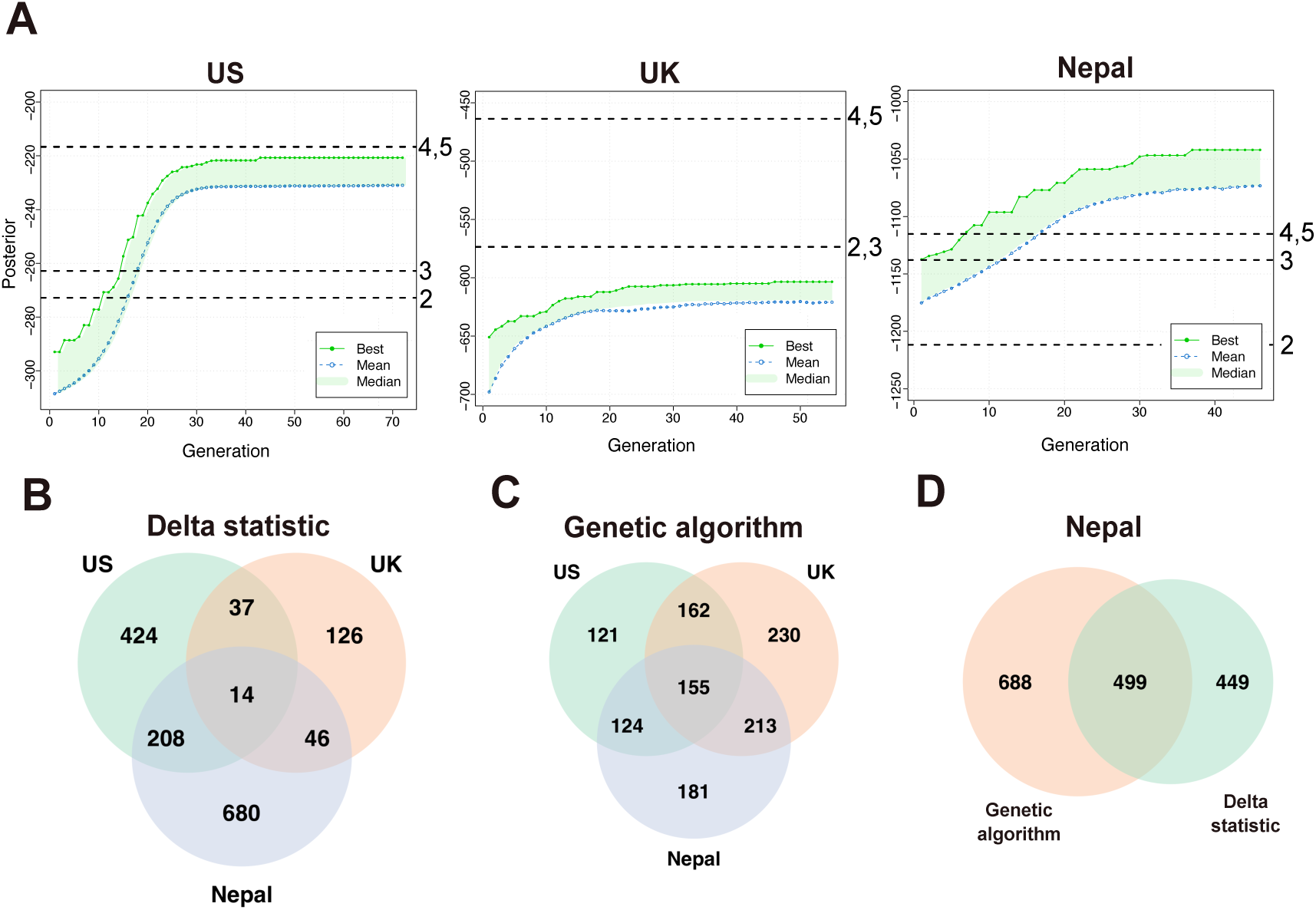
Identifying genes under NFDS using the delta statistic and a genetic algorithm. A) Likelihood of genetic algorithm for the US, UK, and Nepal, and compared to likelihood values of model variants (degrees of freedom 2, 3, 4, and 5) with delta statistic. The x-axis shows iterations of the genetic algorithm (‘generations’). The green shaded area shows the ‘population’ of the genetic algorithm, the green line denotes the best and the blue points denote the average likelihood value. B) Genes under NFDS as determined by delta statistic compared between Nepal, UK, and US. C) Genes under NFDS as determined with genetic algorithm compared between Nepal, UK, and US. D) Genes under NFDS in Nepal, compared between delta statistic and genetic algorithm.

The conceptual difference between the genetic algorithm and the delta statistic is that the genetic algorithm is purely data-informed. The genetic algorithm aims to find the set of genes under NFDS that maximises the likelihood without any constraints. The delta statistic is based on the assumption that genes that change less in frequency are more likely to be under NFDS. This assumption allows ordering the genes by the delta statistic and means that the delta statistic-based *strain-serotype* models simply determine a cut-off for the proportion of genes under NFDS. For now, both approaches depend on datasets spanning pre- and post-vaccine sampling. However, it is plausible that they capture different signals, potentially making one of them less dataset-specific and allowing us to map information from one dataset to another.

We show in Figure 4B that 14 genes were found to be under NFDS in all three countries using the delta statistic. The functional annotation of these genes can be found in Supplementary Table S2. If the number of genes in the overlap is significantly larger than by sampling three subsets of corresponding sizes randomly from the total set of genes, it is likely that the genes in the overlap have some biological relevance; otherwise they are likely not biologically relevant. The expected overlap at random is 23, which is larger than the 14 genes in the NFDS overlap. However, we hypothesised that the delta statistic might be country-specific by design, meaning that small frequency differences might result in a differently ordered gene vector, making it impossible to detect consistent NFDS gene sets.

To test whether the delta statistic hindered us from identifying a consistent NFDS gene set, we applied the genetic algorithm as an alternative method to identify the NFDS gene set. Indeed, the overlap of genes under NFDS between the three countries was 155 genes, many more than in the delta statistic-based analysis; see Figure 4C. However, since the genetic algorithm estimates a higher proportion of genes under NFDS overall, 155 shared genes was still below the expected number of genes under NFDS at random, which was 158.

The overlap between the genes under NFDS using the delta statistic and the genetic algorithm for Nepal is 499, which is slightly larger than the expected 481 genes; see Figure 4C. This suggests that both approaches are somewhat complementary but might detect similar signals.

Overall, our analysis demonstrates that it is difficult to define a consistent NFDS gene set across different countries and vaccination schedules. This highlights the importance of whole genome surveillance to understand vaccine replacement dynamics, since neither serotyping nor targeted sequencing approaches seem to provide sufficient information for the population dynamics model. The lack of consistency in genes predicted to be under NFDS between different locations also emphasizes the need for genomic surveillance in every country of interest, not just in selected ones.

As NFDS is predicted to affect different genes across locations, we sought to characterise the optimal sample size and sampling frequency to inform the population dynamics model and decrease the uncertainty of parameter estimates, ultimately improving our ability to understand the replacement dynamics. This analysis can be interpreted analogously to power analyses that are commonly used to design clinical studies [38, 39].

We simulated the population dynamics model forward with parameter settings based on the US dataset to provide a ground-truth synthetic dataset for comparison. This ground-truth dataset represents the whole *S. pneumoniae* carriage population, rather than a sample. To replicate different sampling strategies, we then sub-sampled the ground-truth dataset by sample size and frequency. In order to compare different sampling strategies, we assumed that there was a fixed and limited ‘sampling budget’, which is represented by the equivalent annual sampling frequency. We assumed that there was a fixed cost per sample and that the sampling frequency did not impact the price of a sample. In other words, a sampling budget of 1% represents 150 annual (1% of 15, 000), 300 biennial, 450 triennial, or 600 quadrennial samples.

We then fitted the population dynamics model to these sub-sampled synthetic datasets and compared the parameter estimates to the true parameter values that were used to generate the simulation; the results of this are shown in Figure 5. First, the vaccination effectiveness and the immigration rate are easier to estimate than the NFDS strength and proportion of genes under NFDS, which can be seen by the fact that the credible intervals tend to be much smaller for the first these parameters. However, the immigration rate estimation appears to be slightly biased towards larger values. Second, for a larger sampling budget (corresponding to sampling 2.5% to 25% annually), sampling frequency has little impact on means and credible interval sizes. Third, and surprisingly, for smaller sampling budgets (especially 0.1% and 0.25% annually) both annual and quadrennial sampling result in biased estimates, whereas biennial and triennial sampling result in better parameter estimates. This might be because the sample size for annual sampling is then very low, lacking the statistical power to detect changes, and the quadrennial sampling might miss critical change points in the data, due to the low sampling frequency.

**Figure 5:**
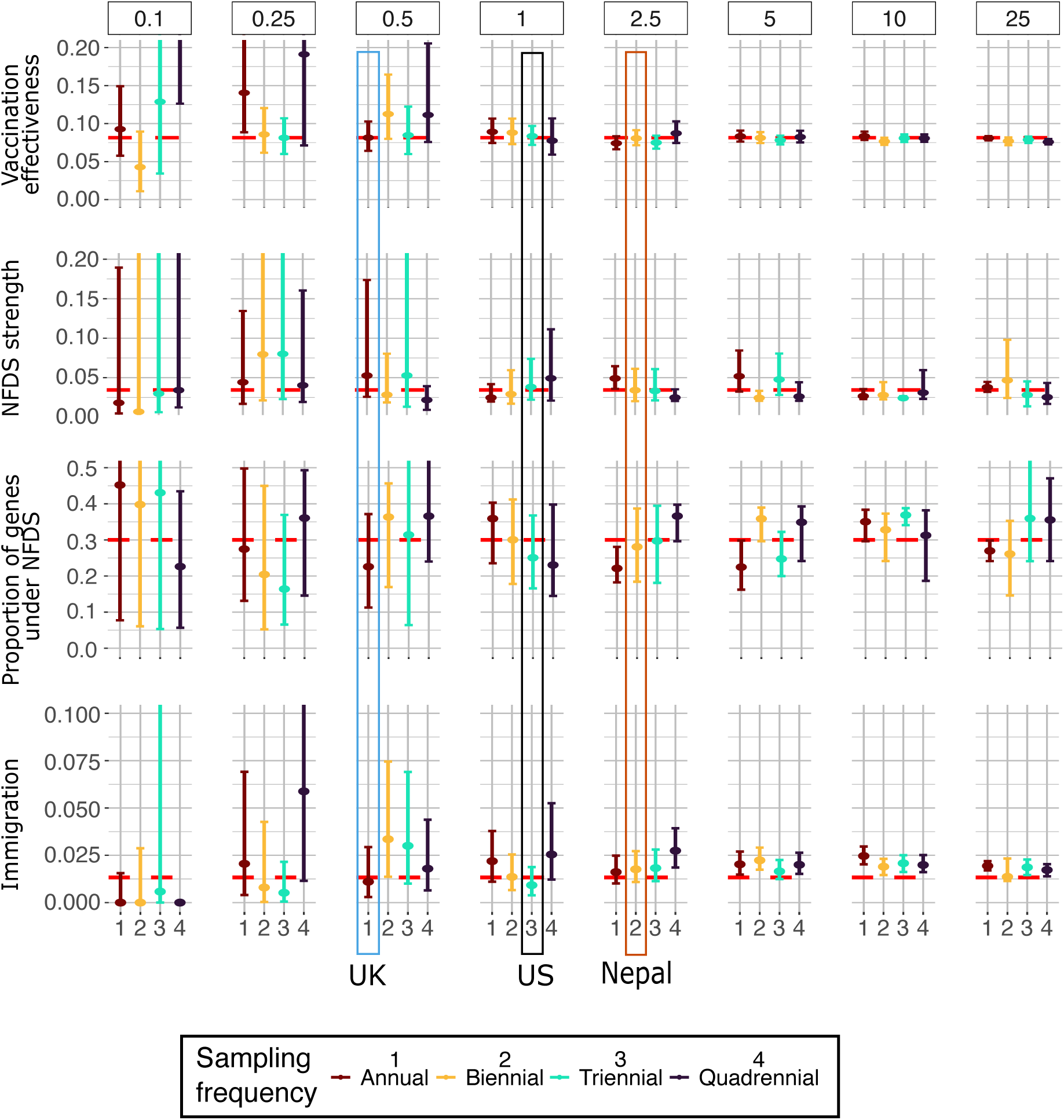
Parameter estimates and credible intervals for fits on synthetic data with different sampling budgets and frequencies. The x-axis and colours show simulated sampling every one, two, three or four years. Y-axis shows parameter values for the four different parameters. Panels show sampling budgets equivalent to sampling 0.1% to 25% annually of the model population of 15, 000 (15 to 3750 annual samples). Original parameter values used to produce synthetic data are shown as red dashed line. Sampling strategies closest to the ones in UK, US, and Nepal are highlighted.

In Figure 5, we also highlighted the sampling strategies that most closely reflect the sampling in the US, UK, and Nepal datasets. Linking back to the parameter estimates for these datasets in Figure 3C, the simulation results suggest that the estimates for the vaccination and immigration parameter can be trusted because the credible intervals of the parameter estimates overlap with the true parameter value and the credible intervals are relatively small. The credible intervals on the NFDS strength and proportion of genes under NFDS however show that the exact parameter estimates should be interpreted with caution because they tend to be larger and mean estimates can be biased. To get a reliable estimate for the NFDS strength and proportion of genes under NFDS, much larger sample sizes are necessary.

Figure 5 is useful for evaluating the impact of the sampling frequency on parameter estimation when assuming a constant sampling budget (per panel). Conversely, for evaluating the impact of the sampling budget when assuming a fixed sampling frequency, grouping the data by sampling frequency can be more informative. Hence, we included Figure S5, which shows the same parameter estimates but grouped by sampling frequency and demonstrates that higher sampling budget tends to result in better parameter estimates and narrower credible intervals.

To answer the question of which sampling strategy results in the best representation of the underlying population dynamics, we used the obtained parameter estimates to simulate the model forward. We then compared those simulations to the ground truth from our synthetic dataset using a cumulative mean squared error (CMSE) across the 20 simulated years. Lower CSME values suggest better agreement between simulation and ground truth. As depicted in Figure 6, the lowest CMSEs were achieved with sampling budgets equivalent to 2.5% annual sampling and higher. Similarly to our results on the parameter estimation, for low sampling budgets of 0.1% and 0.25%, better results are achieved by sampling biennially or triennially, rather than annually or quadrennially. Overall, our findings suggest that when trying to make the most of a limited sampling budget, it would be best to focus sampling efforts on biennial or triennial sampling.

**Figure 6:**
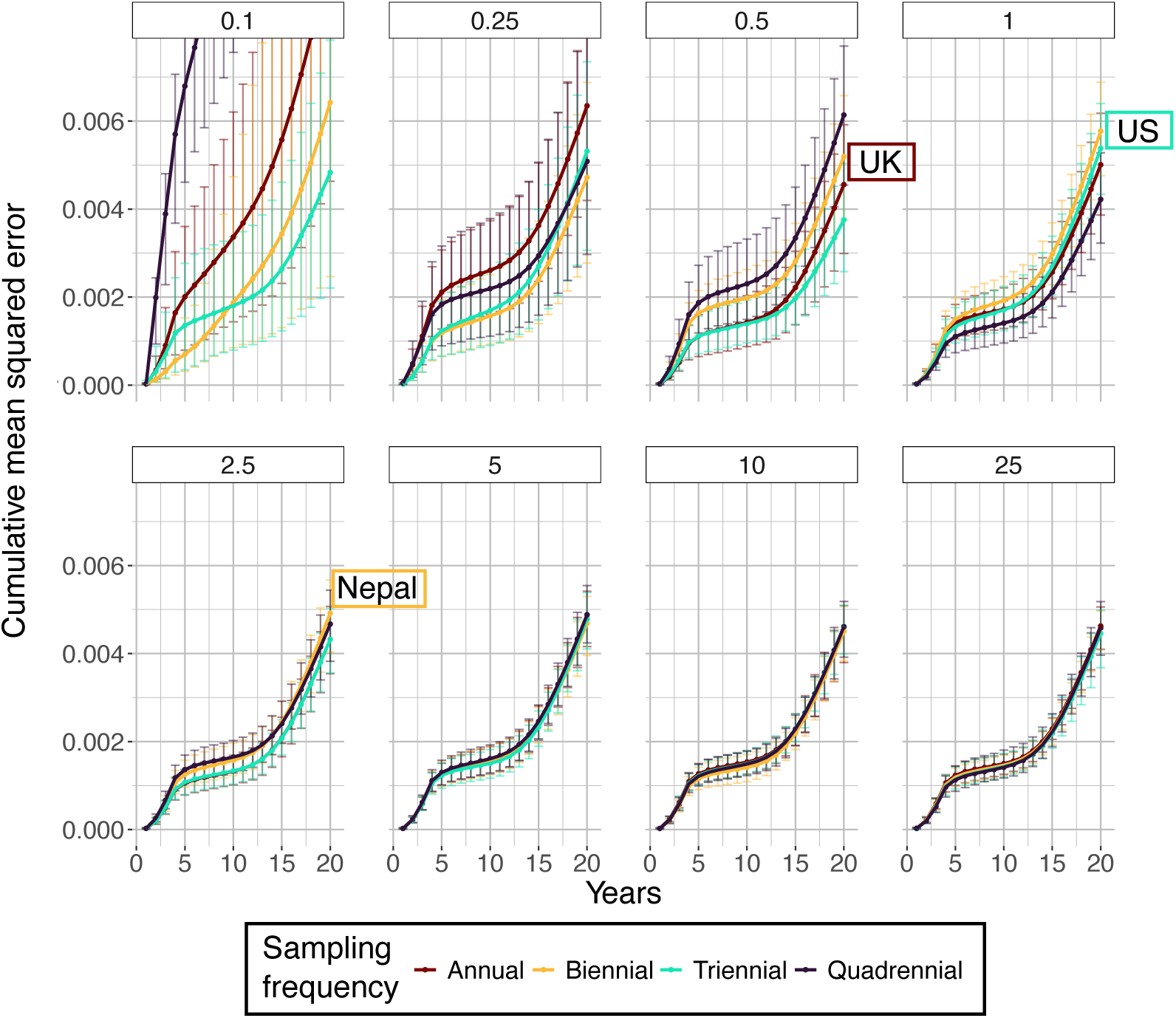
Cumulative mean squared error (CMSE) for model simulation based on fitting parameters on synthetic data with different sample sizes and frequencies. The *x*-axis shows the number of years in the forward simulations. The *y*-axis shows the CMSE. The colours show simulated sampling every one, two, three or four years. The panels show the budget sizes equivalent to annually sampling 0.1 % to 25 % of the model population of 15, 000 (15 to 3750 annual samples). The sample size and frequencies that most closely represent the US, UK, and Nepal dataset are highlighted.

## Discussion

The introduction of PCVs introduced new selection pressures to *S. pneumoniae* populations. This can result in unwanted dynamics, such as the growth of the multi-serotype, multi-drug resistant GPSC10 lineage after the introduction of PCV13 [40]. Mathematical models of population dynamics can help reveal and predict post-vaccine replacement dynamics from genomic surveillance data. These models can also help to inform prospective surveillance strategies. Previous work showed successful application of these models to genomic surveillance data but with limited reusability and without investigating how specific sampling strategies might impact the model’s reliability.

We implemented an optimised population dynamics model incorporating NFDS, which we applied to study post-vaccine replacement dynamics in *S. pneumoniae*. By leveraging the modularisation provided by odin, our model is easier to reuse and adapt, uses reproducible bioinformatics tools, and therefore enables its application to new datasets and regions by public health researchers. We applied our model to genomic surveillance data from the US, UK, and Nepal. Key findings were that the accessory genes inferred to be under NFDS were not significantly similar across countries, and dynamics could not be predicted by serotype alone. This showed how important country-specific whole genome surveillance is and, hence, we applied our model to investigate optimal surveillance strategies. We showed that for a limited sampling budget it is better to sample less frequently — every two or three years rather than every year — to increase the size of each sample. For larger sampling budgets, sampling every year and every two, three or four years gave equivalent results, so the sampling strategy should be entirely informed by practical considerations.

We showed that modern tools can be used to construct a pangenome in an automated manner, suitable for use in NFDS models. However, it is important to note that gene identification and clustering is difficult, especially across multiple datasets, potentially leading to false positive gene identification, which in turn can lead to issues when estimating the impact of NFDS on loci. Consistent pangenomes across different countries are crucial for comparing gene frequencies and genes under NFDS. Yet, creating consistent pangenomes based on existing, country-specific pangenomes is not straightforward, as we showed in the Methods section. A post-hoc mapping of two separate pangenomes with MMseqs2 [25] gave a very different result than a consistent pangenome computed jointly using ggCaller and Panaroo. In the case of our analysis, we opted for computing a consistent pangenome by doing an analysis on all countries jointly but that might not always be feasible, especially due to runtime and memory requirement on large pangenomes. Hence, developing approaches to join existing pangenomes into a consistent one would be desirable.

Our rigorous model selection suggests that NFDS has an effect on many, but probably not all, intermediate-frequency genes. This finding has implications for studying pangenome dynamics, since stable frequencies of some genes might be retained through other means than NFDS, such as balanced rates of gene gain/loss or niches [37, 41, 15] and because some gene frequencies might be less stable than expected under the influence of NFDS.

Microarray techniques, which can be used for serotyping [42], are being extended to detect GPSCs from plate-sweep DNA [43]. It would also be possible to add probes for specific genes under NFDS. This could be a cost-effective alternative to whole genome sequencing for *S. pneumoniae* surveillance, increasing the frequency and sample size to a degree that enables accurate forecasting. However, this would limit the ability to discover new selection effects, and it may be challenging to create universal arrays for all populations. Future work should therefore include investigating the suitability of new microarray data for estimating replacement dynamics using our model.

We investigated how the model accumulates errors in forward simulations using our synthetic dataset but it remains to be investigated how much predictive power the NFDS model has for *S. pneumoniae* post-vaccine dynamics in real datasets. The limited sampling years we had available and the country-specificity of the genes under NFDS did not allow us to investigate this question. However, our model code, which is published in form of an R package, now equips public health surveillance agencies with contemporary data to be able to answer this question in a principled manner.

## Conclusions

We developed a modular, reusable population genetic model that includes parameters for vaccination effectiveness, NFDS, and immigration rates. We showed that our model fits well to genomic carriage data of *S. pneumoniae* from the US, UK, and Nepal. Overall, our work demonstrates how crucial it is to do genomic surveillance of *S. pneumoniae* in every country of interest before and after vaccine introduction. We also discovered that in low-budget settings, it can be beneficial to sample less often in order to have high sample sizes per time point and therefore higher statistical power. Our model is available in the R package STUBENTIGER (**Stu**dying **B**alancing **E**volution (**N**FDS) **T**o **I**nvestigate **Ge**nome **R**eplacement) on GitHub https://github.com/bacpop/Stubentiger.

## Abbreviations

BIC: Bayesian Information Criterion
CMSE: cumulative mean squared error
CPS: capsular polysaccharide
ENA: European Nucleotide Archive
GPS: Global Pneumococcal Sequencing
GPSC: Global Pneumococcal Sequence Cluster
NFDS: Negative frequency-dependent selection
NVT: non-vaccine-type
PCV: pneumococcal conjugate vaccine
VT: vaccine-type

## Ethics approval and consent to participate

Not applicable.

## Consent for publication

Not applicable.

## Availability of data and materials

The whole genome sequences from Kathmandu, Nepal, were obtained from the European Nucleotide Archive (ENA), the accession numbers can be found in Supplementary File 1. Whole genome sequences and metadata for Massachusetts, US [20], and Southampton, UK [21, 22] were previously published.

## Authors’ contributions

J.A.L., L.J.L., C.C., and N.J.C. contributed to conceptualization. L.J.L., J.H., S.T.H., S.W.L., S.S., and A.J.P contributed to data curation. L.J.L. and J.H. contributed to the formal analysis. J.A.L., S.W.L., S.D.B., S.S., and A.J.P. contributed to funding acquisition. L.J.L. contributed to investigation. L.J.L., J.H., J.A.L., C.C., and N.J.C. contributed to methodology. S.W.L., S.D.B., S.S., and A.J.P. contributed to project administration. J.A.L., S.W.L., S.D.B., S.S., and A.J.P. contributed to resources. L.J.L., J.H., S.T.H., M.J.R., and C.C. contributed to software. J.A.L., S.W.L., S.D.B., J.H., C.C., and N.J.C. contributed to supervision. C.C. contributed to validation. J.A.L., L.J.L., C.C., J.H., M.J.R., and S.T.H. contributed to visualization. L.J.L. and J.A.L. contributed to writing – original draft. All authors contributed to writing – review and editing.

## Acknowledgements

The authors thank Víctor Rodríguez Bouza for the discussions around confidence intervals on data.

