## Supplementary Figures and Tables for "A reusable model of pangenome selection informs optimal surveillance strategies over vaccine introductions"

### Supplement

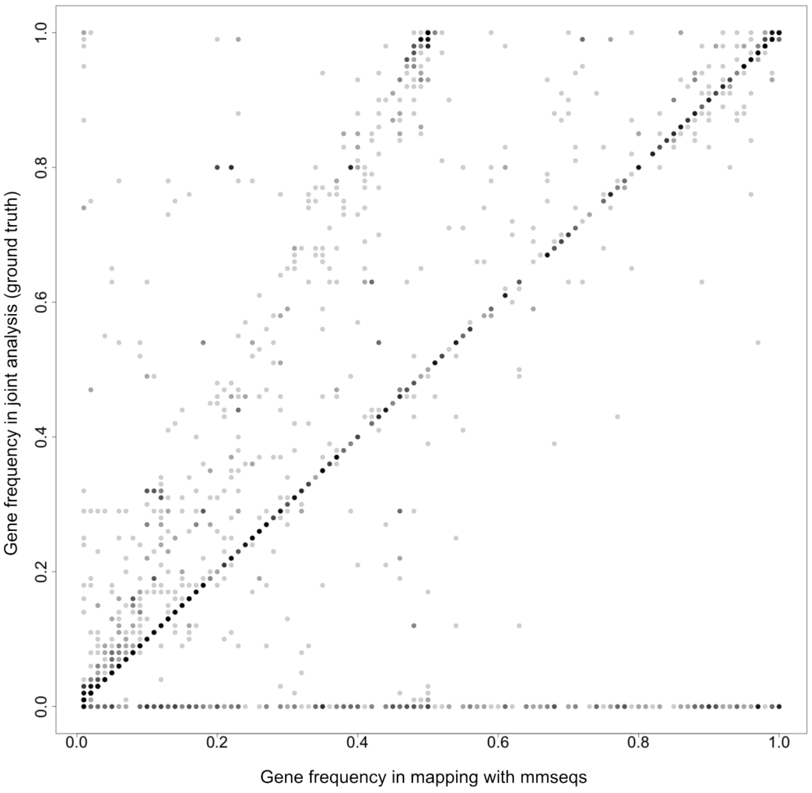


Figure S1: Comparison of post-hoc pangenome mapping using MMseqs and joint pangenome inference using ggCaller and Panaroo.

Table S1: **Comparison of negative log-likelihoods when using COGtriangles or ggCaller/panaroo and Baps or PopPUNK output.** To make likelihood comparable between approaches, it is computed based on the Baps clusters. Best overall likelihood is highlighted in bold. All values based on fits to the US dataset.

| Model | COGtriangles + Baps | ggCaller/panaroo + Baps | COGtriangles + PopPUNK | ggCaller/panaroo + PopPUNK |
|---|---|---|---|---|
| *varying-NFDS model* | −192.88 | −193.10 | −188.60 | **−179.34** |
| *general-NFDS model* | −253.87 | −251.72 | −250.98 | −250.10 |
| *no-NFDS model* | −251.83 | −247.3254 | −260.3223 | −267.9277 |

Table S2: Functional annotation of genes that were found to be under NFDS in US, UK, and Nepal. 14 genes were found to be under NFDS in all countries, out of which 8 had a functional annotation.

| Functional annotation |
|---|
| phosphoenolpyruvate–glycerone phosphotransferase |
| DUF3567 domain-containing protein |
| RelE/StbE replicon stabilization toxin |
| two-peptide bacteriocin subunit BlpN |
| Bacteriocin |
| AzlD domain-containing protein |
| Gp24 |
| Type I restriction-modification system, S subunit |

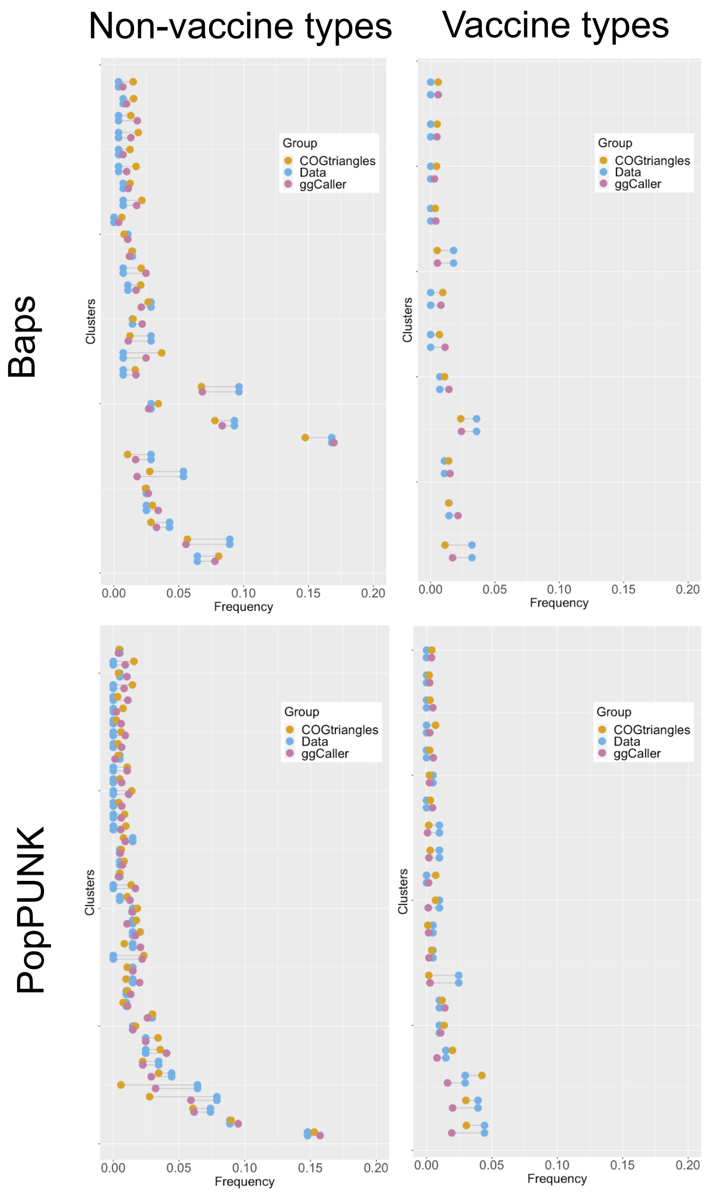


Figure S2: Visual comparison of fits when using COGtriangles or ggCaller/panaroo and Baps or PopPUNK.

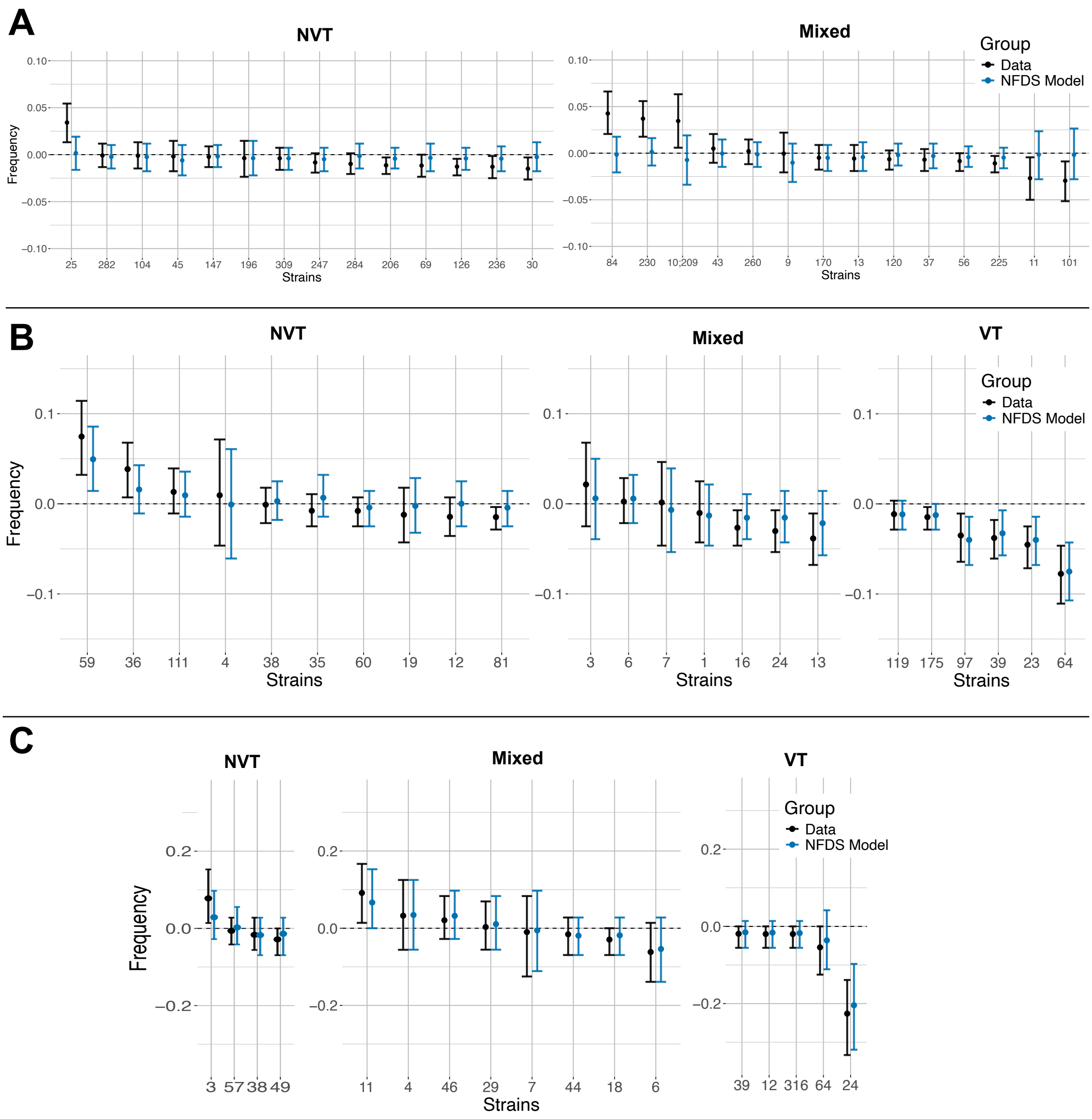


Figure S3: Comparison of GPSC frequency changes in the data and forward simulation of *partial NFDS, strain-serotype* model with mean and 95% confidence intervals. Only serotypes with a frequency of larger than one percent in the pre-vaccine sample are shown. Plots are split into NVT, mixed, and VT GPSCs. Panels show comparison for A) Nepal, B) US, and C) UK.

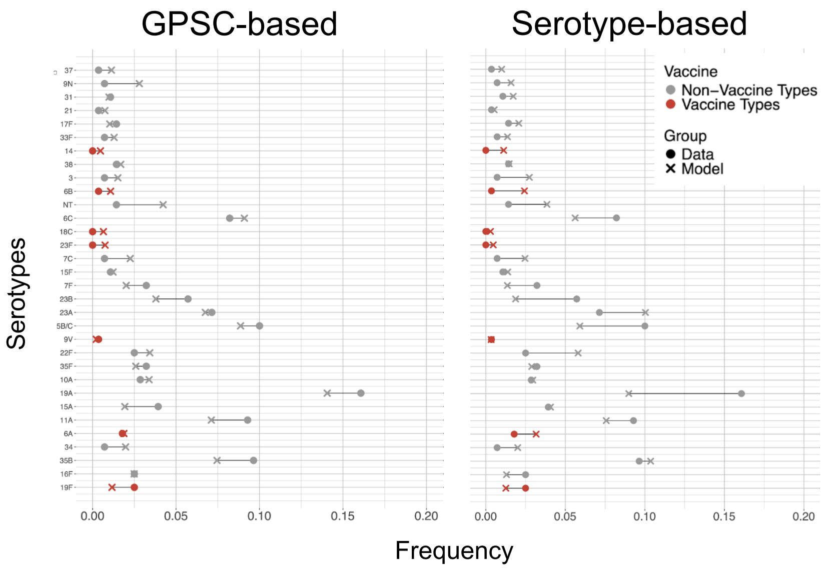


Figure S4: Comparison of GPSC-based model and serotype-based model.

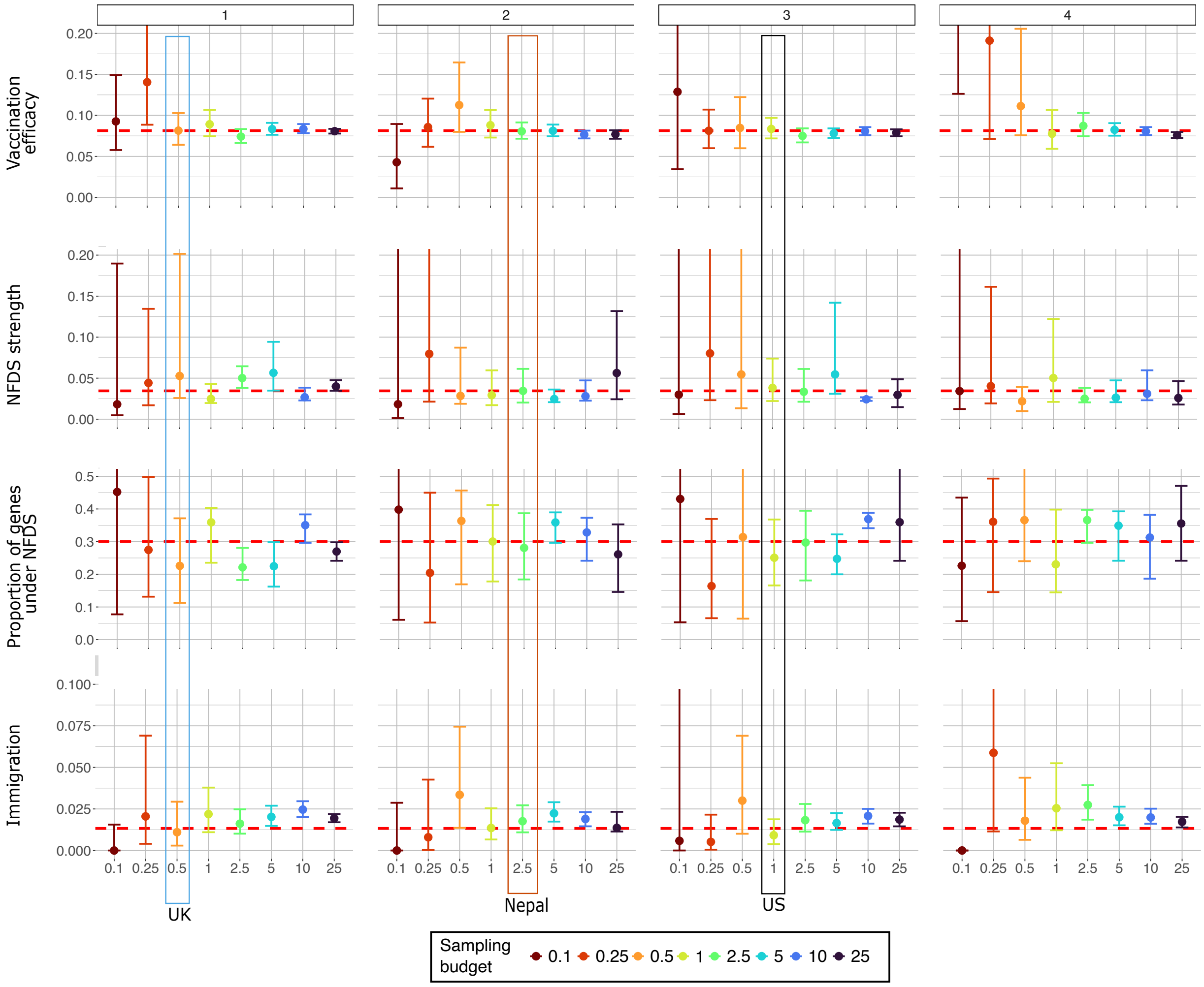


Figure S5: Parameter estimates and credible intervals for fits on synthetic data with different sampling budgets and frequencies. The x-axis and colours show simulated sampling budgets equivalent to sampling 0.1% to 25% annually of the model population of $15,000$. Y-axis shows parameter values for the four different parameters. Panels show sampling every one, two, three or four years. Original parameter values used to produce synthetic data are shown as red dashed line. Sampling strategies closest to the ones in UK, US, and Nepal are highlighted.
